# Global pattern of nitrogen metabolism in marine prokaryotes

**DOI:** 10.1101/2025.10.28.685123

**Authors:** Alexandre Schickele, Hadrien Savioz, Nicolas Gruber, Lionel Guidi, Jean-Olivier Irisson, Meike Vogt

**Author notes:** Correspondence to: Alexandre Schickele.

## Abstract

The ocean nitrogen cycle is driven by an ensemble of metabolic processes sustaining marine ecosystems and ocean productivity. However, the spatial distribution and environmental drivers of its major pathways, i.e., nitrogen fixation, denitrification, assimilatory and dissimilatory nitrate reduction to ammonium (ANRA, DNRA), and nitrification are not well known. Furthermore, the taxonomic composition of the prokaryotes supporting each pathway remain incompletely understood. Leveraging newly assembled global marine metagenomic datasets and a state-of-the-art machine learning framework, we inferred the global biogeography of the genomic potential for key metabolic pathways of the marine nitrogen cycle. This was achieved using a multi-output regression of gene read counts against environmental climatologies. Our results reveal distinct biogeographic patterns of genomic potential: anaerobic or light-inhibited pathways are enriched in high-latitude regions, eastern boundary upwelling systems, and deeper ocean layers, while nitrogen fixation and ANRA dominate in oligotrophic gyres. These patterns are consistent with known metabolic strategies, model-based estimates, and underlying taxonomy. Indeed, we identify that *Cyanobacteria* associate primarily with aerobic, biosynthetic pathways, while *Gammaproteobacteria* and *Nitrososphaeria* encode for nitrogen transformations related to energy requirements. By coupling microbial community composition with genome-level information, our approach advances understanding of the microbial foundations of nitrogen transformation pathways and offers new insights on underrepresented processes into biogeochemical models. We highlight the growing value of omic data to better understand marine ecosystem function in relation to environmental gradients and community composition, and their use as a potential observation-based alternative or complement to biogeochemical models.

## 1. Introduction

As one of the chemically most versatile elements, nitrogen exists in the ocean in a myriad of forms, with the vast majority of chemical transformations between these forms being undertaken by marine organisms, especially prokaryotes (Herrero et al., 2019; Hutchins and Capone, 2022; Zehr and Kudela, 2011). Two key metabolic requirements drive these transformations. The first metabolic requirement is a consequence of nitrogen being an essential element for life, as it is required to form amino-acids and other key cellular component such as nucleotides. To achieve this, nitrogen needs to be taken up from the environment and brought into its most reduced oxidation state (N^-III^), required for most organic forms of nitrogen (Herrero et al., 2019). The second group of nitrogen transformations is driven by the need of organisms for energy, through either autotrophic or heterotrophic forms of metabolisms. The organisms thereby take advantage of the wide range of nitrogen’s oxidation states (-III to +V), providing opportunities for a large set of exothermic redox processes.

One of the fascinating and at the same time puzzling aspects of the marine nitrogen cycle is how these two groups of metabolic processes interdepend on each other. This is especially relevant when considering that most nitrogen in the ocean exists in the form of dinitrogen gas (N_2_), which is not directly bioavailable and largely inert (Herrero et al., 2019). In fact, the total oceanic inventory of N₂ is more than 10 times higher than the sum of nitrate (NO₃⁻), nitrite (NO₂⁻) and ammonium (NH_4_^+^), which constitute the primary forms of biologically available nitrogen and are often referred to as “fixed nitrogen” (Gruber, 2008). Furthermore, nitrate is an order of magnitude more abundant in the surface ocean than ammonium, even though the latter is the preferred source of N for uptake. The dearth of ammonium is a consequence of marine prokaryotes oxidizing ammonium to nitrate as part of their chemo-autotrophic metabolism (Ward, 2008). Also, many prokaryotes use nitrate as a terminal electron acceptor, thereby converting it to biounavailable N_2_. Thus, the energy-driven metabolic transformations tend to affect the availability of fixed nitrogen for sustaining life and productivity in the ocean. This affects in turn the ocean carbon cycle and climate (Gruber, 2004), given the key role of fixed nitrogen as a factor limiting primary and export production across vast ocean areas (Moore et al., 2013).

Two very different transformation processes enable the first metabolic requirement, i.e., the uptake of nitrogen from the environment to build cellular constituents. The first one is the Assimilatory Nitrate Reduction to Ammonium (ANRA; **Fig. 1**, pathway 1)(Zhang et al., 2020), whereby nitrate is taken up by the cell and intercellularly reduced to the oxidation state of ammonium. Nearly all phototrophic organisms living in the ocean have the corresponding enzymes, making this metabolic process ubiquitous. Its magnitude is relatively tightly coupled to the ocean’s primary production, 10 to 20% of which is supported by ANRA (Gruber, 2008; Zhang et al., 2020). This yields an annual rate of about 850 to 1700 Tg N yr^-1^ (Gruber, 2008). The second metabolic transformation supporting the uptake of nitrogen from the environment is nitrogen fixation (**Fig. 1**, pathway 2). Here N₂ is reduced by nitrogenase, to ammonium. This process is energetically costly and only undertaken by a relatively small group of microbes named diazotrophs (Carpenter and Capone, 2008). The total magnitude of nitrogen fixation in the marine environment is of the order of 100 to 200 Tg N yr^-1^, thus an order of magnitude smaller than ANRA (Wang et al., 2019). While most of marine nitrogen fixation is undertaken by photoautotrophic organisms in the low latitudes and in waters that are low in nitrate (Wang et al., 2019), it has also been observed in high latitudes, in waters rich in nitrate (Von Friesen and Riemann 2020)(Von Friesen and Riemann, 2020), and also associated with heterotrophy (Bombar et al., 2016; Von Friesen and Riemann, 2020; Zehr and Capone, 2020).

**Figure 1:**
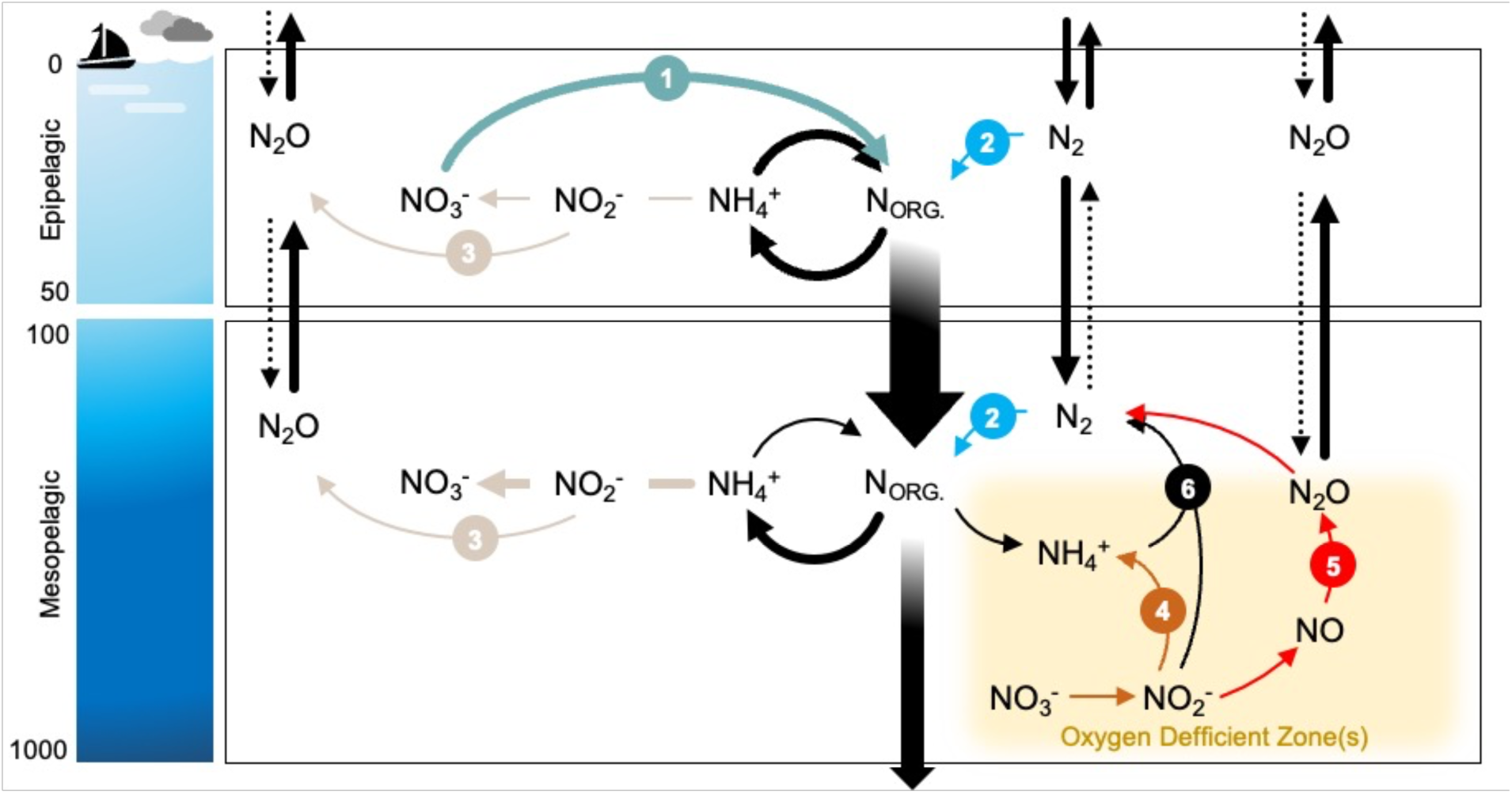
The marine nitrogen cycle as considered in our study. The diagram represents the chemical transformations and essential metabolic pathways of the marine nitrogen cycle compartmented across atmosphere, epipelagic, mesopelagic and oxygen deficient zones. The metabolic pathways performed by marine prokaryotes and considered in our study are indicated by coloured arrows and numbers. Those include: (1) Assimilatory nitrate reduction to ammonium (ANRA); (2) Nitrogen fixation; (3) Nitrification; (4) Dissimilatory nitrate reduction to ammonium (DNRA); (5) Denitrification. Black arrows represent pathways not considered in this study such as (6) Anaerobic ammonia oxidation (Anammox); and physical transport processes of nitrogenous compounds.

The transformations driven by the metabolic requirement for energy can be differentiated into those associated with autotrophy and those with heterotrophy. Furthermore, the free enthalpy associated with the redox reactions involving nitrogen are highly dependent on the presence or absence of oxygen, such that completely different sets of metabolic transformations dominate under oxic or anoxic conditions (Lam and Kuypers, 2011; Zhang et al., 2023). Under oxic conditions, a group of chemo-autotrophic organisms oxidize ammonium to nitrate in two distinct steps, jointly referred to as nitrification. In the first step ammonium-oxidizing bacteria and archaea convert ammonium to nitrite, followed by nitrite-oxidizing bacteria oxidizing nitrite to nitrate (**Fig. 1**, pathway 3) (Ward, 2008; Zhang et al., 2020). These two processes tend to be inhibited by light, such that their prevalence in the upper layers is relatively low. But nitrification thrives in the layers just below that, in the depth range between 100 and 500 m in the mesopelagic zone, where it oxidizes the ammonium that is being released from the organic nitrogen that is sinking or transported downwards from the near-surface layers. This leads nitrification to turn around about the same magnitude of nitrogen as ANRA, that is about 1000 Tg N yr^-1^. In anoxic environments, such as oxygen deficient zones (ODZs), oxidized nitrogen compounds serve as electron acceptors for anaerobic respiration. In Dissimilatory Nitrate Reduction to Ammonium (DNRA; **Fig. 1**, pathway 4), nitrate is sequentially reduced to nitrite and then to ammonium, with energy conservation linked to cellular respiration (Wang et al., 2024). Denitrification reduces nitrate to N₂ in four steps, generating nitric oxide (NO) and nitrous oxide (N₂O) as intermediates (**Fig. 1**, pathway 5). In the water column, denitrification is believed to be quantitatively much more important than DNRA, globally reducing around 65 Tg N yr-1 (Battaglia and Joos, 2018). This is still more than a magnitude smaller than nitrification. Being an obligatory anaerobic process, denitrification is confined to the Arabian Sea, the eastern tropical Pacific, and a small portion of the eastern (sub)tropical North and South Atlantic. Finally, some prokaryotes perform anaerobic ammonium oxidation (Anammox; **Fig. 1**, pathway 6), which consists in oxidizing ammonium using nitrite as an electron acceptor, resulting in the production of N_2_ (Rios-Del Toro and Cervantes, 2019). While knowledge on this complex ensemble of metabolic processes associated with the nitrogen cycle has grown rapidly in recent decades (Hutchins and Capone, 2022), our knowledge of the global pattern of these processes remains rather limited (Zhang et al., 2020). Moreover, the microbial communities associated with these processes and their niche partitioning also remain insufficiently understood, and they are poorly represented in culture collections (Hugenholtz, 2002).

Over the past decade, large-scale marine biological surveys have expanded their global coverage and incorporated advanced observation techniques, including metagenomic sequencing (Sunagawa et al., 2015). Genome-resolved metagenomics from programs such as Tara Oceans and BIOGEOTRACES have enabled the reconstruction of approximately 35,000 genomes (Paoli et al., 2022). These genomes serve as taxonomic units that are functionally annotated based on the enzymes they encode and the metabolic processes these enzymes support. The relative abundance of such enzymes, referred to as the genomic potential for a given metabolic process, can be quantified using genome-wide metagenomic read coverage, offering a direct link between microbial identity and function (Sunagawa et al., 2015). Building on these data, recent studies have mapped the global distribution of processes related to energetic metabolism (Salazar et al., 2019) and carbon concentration mechanisms associated with photosynthesis (Schickele et al., 2024). Similarly, metagenomics *in-situ* observations may provide valuable insights into genes involved in nitrogen transformations and the microbial communities that encode them – including the one associated with ANRA, DNRA, and nitrification – which remain underrepresented in most biogeochemical models (Wang et al., 2024; Zhang et al., 2020). This makes metagenomics particularly promising to study how the genomic potential supporting nitrogen transformations is distributed in the global ocean, which microbial communities mediate these processes, and under which environmental conditions they occur. However, marine biological sampling remains spatially and temporally limited, especially in remote ocean regions, providing only a fragmented view of the global distribution of the genomic potential for nitrogen transformations (e.g., Salazar et al., 2019).

Habitat modelling addresses this limitation. Rooted in ecological niche theory, it is widely used to predict the spatial distribution of organisms based on the environmental conditions at observed locations (Peterson and Soberón, 2012). By statistically relating occurrences to environmental conditions, habitat models enable the generation of spatially continuous distribution estimates from sparse *in-situ* observations. Therefore, they provide a continuous biogeography of taxon and the functional traits they are associated with (Peterson and Soberón, 2012). In the marine environment – where direct sampling is limited and uneven in space and time – this approach has been particularly valuable for inferring macroecological patterns (e.g., Beaugrand et al., 2019). Marine microorganisms, including prokaryotes and the biogeochemical transformations they mediate, are especially sensitive to environmental gradients that shape their physiology, metabolic strategies, and ecological functions (Hutchins and Fu, 2017). Recent advances in genome-resolved metagenomics, combined with machine learning, now allow habitat modelling to move beyond species presence toward predicting the distribution of genomic potential for key biogeochemical processes encoded in microbial DNA (Delmont et al., 2022; Paoli et al., 2022; Schickele et al., 2024).

Building on recent efforts to map the distribution of marine microbial communities and their encoded functions (e.g., Coelho et al., 2022; Richter et al., 2022; Schickele et al., 2024), we investigate whether genome-resolved metagenomics can provide ecologically and biogeochemically meaningful insights into the spatial distribution of nitrogen transformations in the global ocean. We hypothesize that genome-resolved metagenomic data can distinguish between different types of nitrogen transformations based on their metabolic requirements and their association with environmental context, such as oxygen availability. Specifically, we ask whether metagenomics can (i) capture distinct spatial patterns between the genomic potential for nitrogen transformations related to cellular biosynthesis and electron transport, (ii) reveal distinct environmental conditions and microbial communities driving these two types of transformations, and (iii) serve as an observation-based complementary or alternative to conventional biogeochemical measurements for monitoring marine nitrogen cycling in space and time. To address these questions, we use the CEPHALOPOD habitat modelling framework (Schickele et al., 2025) that enables the simultaneous projection of metabolic processes encoded in microbial genomes, accounts for interactions between these processes, the environmental conditions they occur in, and the microbial communities they are associated with. We apply this to project the distributions of the genomic potential for five key nitrogen transformation pathways – nitrogen fixation, ANRA, nitrification, DNRA, and denitrification – across the epipelagic and mesopelagic layers of the global ocean. Doing so, explore not only where the genomic potential for each nitrogen transformation is distributed, but also how different pathways co-occur, or partition across environmental conditions, offering a new perspective on the functional organization of nitrogen cycling in marine microbial communities.

## 2. Methods

### 2.1. Data source and selection

#### 2.1.1. Metagenomic data collection

We investigated how the genomic potential supporting nitrogen transformation distribute in the global ocean using a comprehensive collection of c.a. 35,000 prokaryotic genomes derived from multiple global ocean expeditions, including – but not limited to – Tara Oceans and BIOGEOTRACES expeditions (Paoli et al., 2022). This dataset comprises approximately 10,000 genomes from cultivated isolates and single cell amplified genomes, along with approximately 26,000 predominantly bacterial and archaeal Metagenome-Assembled Genomes (MAGs). These genomes are annotated taxonomically (i.e., from domain to species level) and functionally (i.e., metabolic pathway and enzyme level; according to the Kyoto Encyclopedia of Genes and Genomes; KEGG) (Kanehisa and Goto, 2000). They are also associated with georeferenced information from 1,038 seawater samples collected at 215 globally distributed stations, monthly resolved (**Fig. 2a**), spanning depth layers from the surface ocean to 5,600 meters, and encompassing the epipelagic, mesopelagic, and bathypelagic zones (Paoli et al., 2022). The collection includes prokaryote-enriched size fractions ranging from 0.2 to 3 μm, providing broad spatial and ecological coverage from which we retrieved taxonomic and functional information related to nitrogen transformation pathways in prokaryotes.

**Figure 2:**
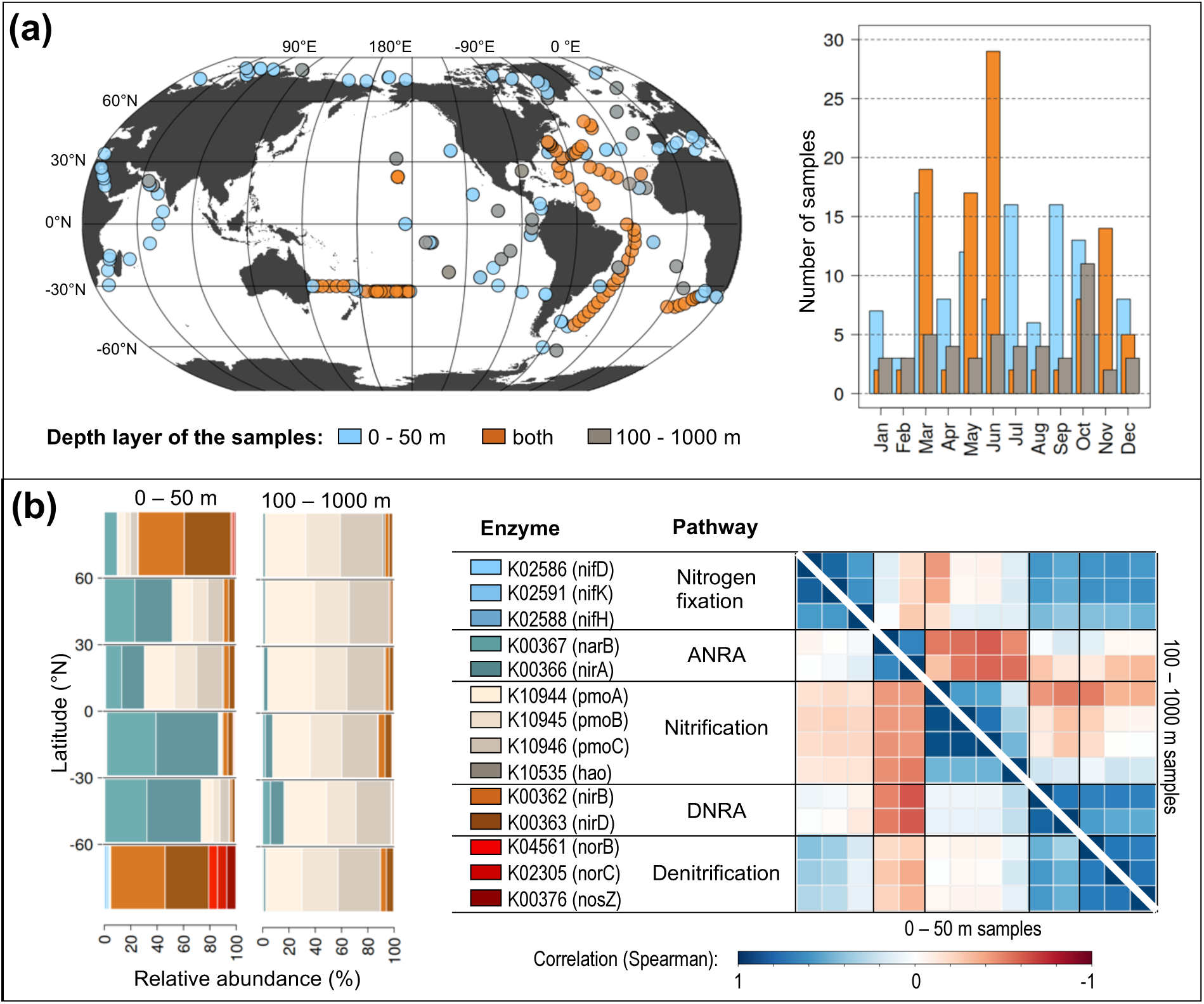
Overview of the metagenomic data considered with, (a) their observed distribution in space, time, and depth and (b) their functional annotations, relative abundance latitudinal profile and Spearman correlation at the KEGG Orthologs (KO) enzyme and metabolic pathway-level.

#### 2.1.2. Selection of enzymes supporting nitrogen transformations

To extract the genomic potential associated with each nitrogen transformation pathway, we first filtered genomes based on their gene functional annotations using two criteria: (1) the gene must encode for an enzyme specific to one of the targeted nitrogen transformation pathways, and (2) the gene must be present in more than 50 individual observations to ensure sufficient spatial coverage to estimate its climatological distribution (i.e., see CEPHALOPOD benchmark in Schickele et al., 2025). To exclude viral observations, we further restricted our analysis to genomes from size fractions corresponding to free-living prokaryotes: 0.22 – 3 μm, 0.22 – 1.6 μm, and >0.2 μm. Finally, we associated these observations with depth layers of 0 – 50 m and 100 – 1,000 m, here considered as the epipelagic (416 samples) and mesopelagic (319 samples) layers of the ocean, respectively. We excluded samples collected between 50 and 100 m depth (173 samples; 16.6%) because this depth interval frequently intersects the mixed layer (Boyer et al., 2018). At a climatological scale, this prevents a robust assignment of samples to either mixed-layer or sub–mixed-layer regimes. This filtering resulted in a set of 14 genes encoding enzymes specific to nitrogen fixation, ANRA, nitrification, DNRA, and denitrification, based on the KEGG classification (**Fig. 2b**)(Kanehisa and Goto, 2000). Among the three enzymes associated with the Anammox pathway (**Fig. 1**, pathway 6) and found in our metagenomic data collection (Paoli et al., 2022), Nitrite reductase NO forming enzymes *K00368* and *K15864* did not meet the specificity criteria as they are shared with Denitrification, and Hydrazine synthase *K20934* was present in less than 50 observations. Therefore, we did not consider the Anammox pathway (**Fig. 1**, pathway 6) in subsequent modelling steps. The quality of the metagenomic data was evaluated using both the ‘lineage workflow’ of CheckM (v1.0.13) and Anvi’o (v5.5.0) by means of a completeness and contamination metric which represents the fraction of marker genes of the prokaryotic clade respectively included or miss-assigned in the MAGs (Paoli et al., 2022). The mean (*±* standard deviation) completeness (90 *± 16* %) and contamination (1.6 *± 1.7* %) across all enzymes encoding for nitrogen transformation processes align with ‘good quality’ according to community standards (completeness ≥ 90 % and contamination ≤ 5 %; Bowers et al., 2017). To evaluate whether nitrogen transformation pathways could be modelled as unified biological processes rather than as individual enzymes, we conducted a Spearman correlation analysis among enzyme-level relative abundances within each pathway, separately for the epipelagic and mesopelagic layers (**Fig. 2b**). Enzyme relative abundances within each pathway were highly correlated, with median Spearman coefficients exceeding 0.80 in the epipelagic and 0.82 in the mesopelagic layer – except for nitrification in the mesopelagic layer, which showed a lower correlation (0.64; **Fig. 2b**). Based on this consistency, we aggregated enzyme-level signals to the pathway-level by averaging the relative abundances of genes encoding enzymes within each KEGG-defined nitrogen transformation pathway (Kanehisa and Goto, 2000), and used these aggregated values as the unit of analysis in subsequent modeling.

#### 2.1.3. Environmental feature collection

Habitat modelling relates biological observations to the environmental conditions in which they occur (Peterson and Soberón, 2012). To capture the environmental drivers of gene distribution associated with nitrogen transformation pathways, we selected 47 monthly resolved, observation-based environmental climatologies (i.e., hereafter named features) with a spatial resolution of 1° x 1°, at the global scale (**Table S1**) (Schickele et al., 2025). These features describe the physical (e.g., sea surface temperature), chemical (e.g., nutrient concentrations, carbonate chemistry), and biological (e.g., chlorophyll-a concentration, primary production) properties of water masses, as well as their circulation and turbulence characteristics (e.g., eddy kinetic energy, EKE). At climatological scales, EKE does not aim to assess sub-mesoscale processes, but rather highlight large scale ocean currents (e.g., Gulf Stream, Alghulas current). When applicable (i.e., excluding satellite-derived products), environmental fields were integrated over the depth ranges of biological sampling: 0–50 m for the epipelagic layer and 100–1000 m for the mesopelagic layer.

### 2.2. Modelling workflow

To extract the biogeographic patterns of nitrogen transformation pathways from metagenomic data, we used CEPHALOPOD, a recently developed framework that standardizes habitat modelling across diverse marine datasets, including those with relative abundances of metagenomic reads (Schickele et al., 2025). This framework uses an ensemble approach of statistical and machine learning methods, a parsimonious environmental features selection and an exhaustive uncertainty assessment to extract patterns in biological information, and to provide projections to the global scale.

#### 2.2.1. Pre-processing

In the pre-processing step, environmental features are first matched in space and time with the biological data to minimize sampling bias using a common, monthly 1x1° grid. To this end, biological target values are averaged within each spatio-temporal grid cell and integrated across the corresponding depth layer. To ensure robust model inputs, we applied a three-step pre-processing procedure. First, outliers in the biological inputs were removed using a Z-score threshold of ±2.5 standard deviations from the mean target value (Knecht et al., 2023). Second, environmental features unlikely to explain target variability were excluded: CEPHALOPOD filters out features that exhibit stronger Spearman correlation or mutual information with a null target (i.e., randomly sampled values within the target range) than with the actual biological target (Schickele et al., 2025). Third, multicollinearity was addressed by computing pairwise Pearson correlations among features at target locations (**Fig. S1**). Within each group of collinear features (r > 0.8), only the feature with the strongest association (via correlation or mutual information) with the biological target was retained (Schickele et al., 2025). Finally, to identify the regions where SDM extrapolates outside environmental conditions of observations, CEPHALOPOD performs a Multivariate Environmental Similarity Surfaces (MESS; Elith et al., 2010).

#### 2.2.2. Algorithm training, evaluation, and projection

For proportional data, CEPHALOPOD employs a Multivariate Boosted Tree Regressor (MBTR), a recently developed algorithm designed specifically for multivariate response variables (Nespoli and Medici, 2022; Schickele et al., 2024). The algorithm training follows a *n*-fold cross-validation procedure (here *n = 5*), where the dataset is split into *n* subsets: each model is trained *n* times on *n-1* folds, with the remaining fold reserved for testing (Araújo et al., 2019). Hyperparameters for the MBTR algorithm are first optimized across the *n*-training folds. Using these optimal settings, CEPHALOPOD evaluates predictive performance by applying the algorithm to predict the biological targets to environmental conditions corresponding to the *n*-test sets. To assess uncertainty in spatial projections, a bootstrap resampling approach is performed: CEPHALOPOD generates 100 bootstrap replicates by sampling the original dataset with replacement, maintaining the same sample size. For each replicate, a full MBTR algorithm is trained using the previously determined best hyperparameters. The trained algorithm is then used to project monthly genomic potential for nitrogen transformation pathways globally, based on the environmental features associated with each grid cell (Schickele et al., 2025).

#### 2.2.3. Final quality assessment

CEPHALOPOD incorporates a multi-criteria quality control system specifically tailored for multivariate species distribution models (SDMs), integrating four standardized quality checks to ensure robustness and interpretability of model outputs. First, the *a priori* relevance of the selected environmental features is evaluated by comparing their average Spearman correlation and mutual information with the biological target against a null target (i.e., randomly permuted target values). To pass this check, the combined signal strength must exceed that of the null target by at least 0.05, following an adaptation of Knecht et al. (2023) and Kinney & Atwal (2014). Second, CEPHALOPOD assesses predictive performance using the evaluation folds from the cross-validation. For proportional (i.e., multivariate) data, model skill is quantified using a multivariate R² (Pearson), ranging from –1 to 1, and estimating the compositional prediction accuracy (i.e., as the model optimises a compositional loss function; Nespoli and Medici, 2022). A minimum threshold of 0.25 is required to ensure sufficient predictive power (Schickele et al., 2025). Third, the relative importance of features is used to evaluate algorithm interpretability using the tidymodels R library (v1.1.1). To ensure ecological interpretability, MBTR is only retained if the three most influential features collectively explain at least 50% of the total variable importance (see **Text S1**, Sect. 1) (Schickele et al., 2025). Due to the multivariate nature of MBTR, the variable importance at the algorithm level is associated to the target composition, the variable importance associated to each target was estimated through a Redundancy Analysis (RDA; see **Text S1**, Sect. 2). Fourth, CEPHALOPOD quantifies projection uncertainty using the Normalized Standard Deviation (NSD), defined as the standard deviation across bootstrap replicates normalized by the mean value across all geographical cells. MBTR is retained only if the global average NSD remains below 0.5. We require MBTR to pass all four quality checks to be used for generating global genomic potential for nitrogen transformation pathways maps, ensuring both predictive performance and interpretability in multivariate modelling context (Schickele et al., 2025).

## 3. Results

### 3.1. Model quality assessment

Our spatial projections of the genomic potential for nitrogen transformation pathways successfully passed all CEPHALOPOD quality checks (see sect. 2.2.3.). We estimated a predictive performance (R2 > 0.25) of 0.44 and 0.42 for the epi- and mesopelagic models, respectively. The three features explaining most of the variance in the observation presented a cumulative feature importance (> 50 %) of 75 and 81 % for the epi- and mesopelagic models, respectively. Finally, we estimated a Normalized Standard Deviation of the projections across bootstraps resampling (NSD < 0.5) of 0.24 and 0.21 for the epi- and mesopelagic models, respectively. This means that our spatial projections show confidence in reproducing the observed distribution of genomic potential (Schickele et al., 2025). We highlight however that projections in high latitudes and coastal areas are subject to extrapolation outside the environmental conditions of the observations and should therefore be considered with caution (**Fig. S2**).

### 3.2. Dominance patterns in nitrogen transformation pathways

Here, we present the composition in nitrogen transformation pathways across open-ocean biomes, for both epi- and mesopelagic layers (**Text S1**, **Fig. 3**). This composition is calculated from the yearly averaged projection of relative abundance of metagenomic reads associated to each pathway and should be interpreted as the genomic potential for each pathway.

**Figure 3:**
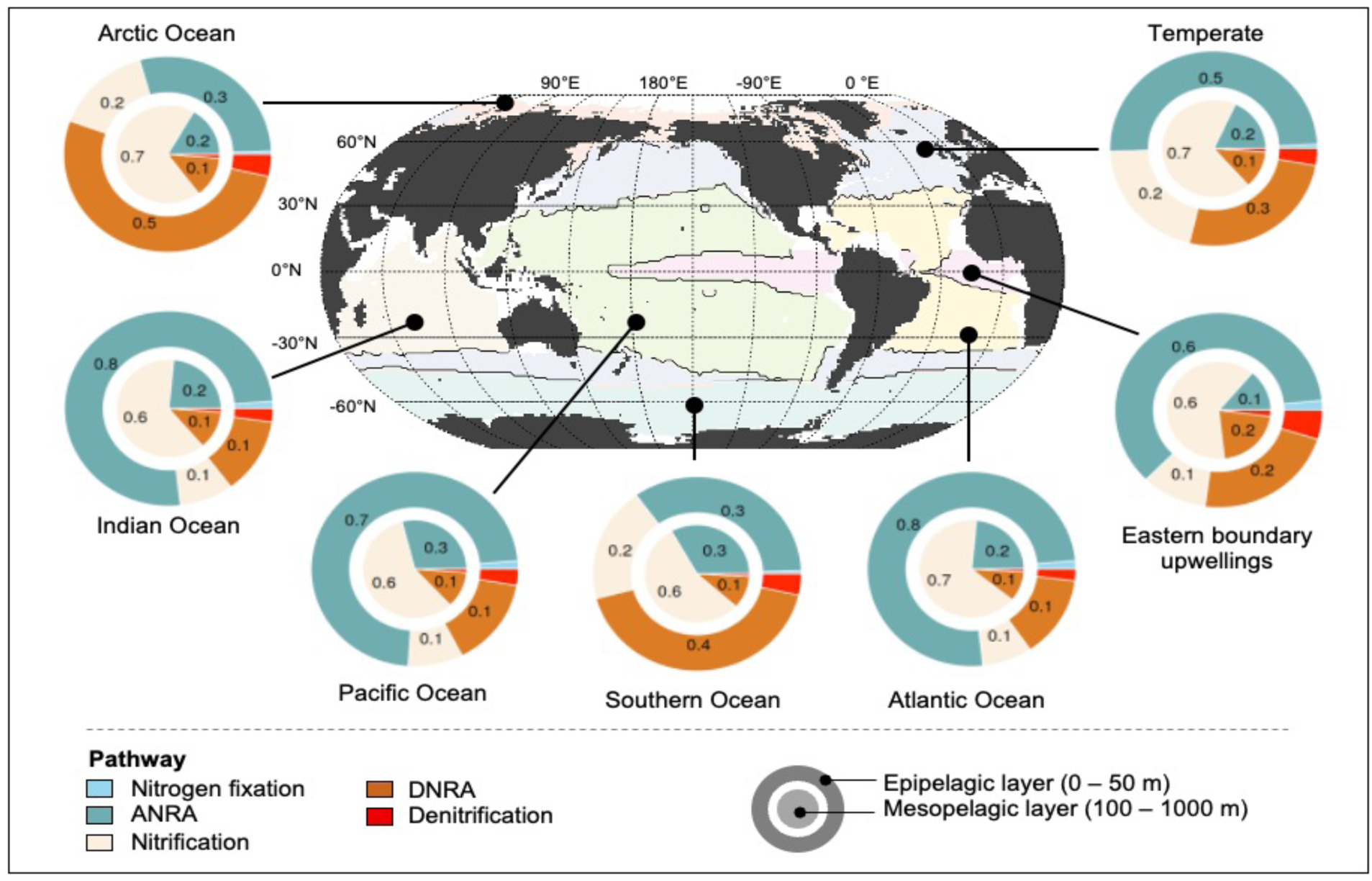
Overview of the nitrogen transformation pathway distribution in the epi- and mesopelagic layers of the global ocean, aggregated by open-ocean biomes adapted from Fay and McKinley (2014). Each biome is associated to a single colour for readability. Pie charts represent the average genomic potential for each nitrogen transformation pathway. The epi- and mesopelagic layers are represented in the outer and inner pie charts, respectively.

First, we observe a clear dominance of nitrogen transformation pathways associated with bioavailable over bio-unavailable forms of nitrogen – namely ANRA, DNRA, and nitrification – in both epi- and mesopelagic layers (**Fig. 3**). In the epipelagic layer, ANRA accounts for the largest share of the genomic potential (0.59), followed by DNRA (0.23) and nitrification (0.13). In contrast, pathways associated with bio-unavailable nitrogen, such as denitrification (0.03) and nitrogen fixation (0.01), are comparatively minor (**Fig. 3**, inner pie charts). A similar pattern is observed in the mesopelagic layer, where nitrification emerges as the dominant pathway (0.62), followed by ANRA (0.25) and DNRA (0.11). Again, denitrification (0.01) and nitrogen fixation (<0.01) represent a small fraction of the total genomic potential (**Fig. 3**, inner pie charts). These results suggest that the genomic potential associated with bioavailable nitrogen transformation is more widespread and genomically encoded than for pathways involving bio-unavailable nitrogen forms, especially in deeper ocean layers. This also means that the use of nitrogen transformation for cellular biosynthesis or energetic requirements is not the first order gradient spatially, but the association with bioavailable or bio-unavailable forms of nitrogen. Due to this dominance pattern, subsequent analyses will therefore be separated between the bioavailable and bio-unavailable nitrogen transformations (*sect. 3.3. and 3.4*). Second, we identify a pronounced latitudinal gradient in the projected genomic potential associated with DNRA in the epipelagic layer (**Fig. 3**), as shown in the *in-situ* observations (**Fig. 2b**). Indeed, DNRA represents a substantial proportion of nitrogen transformation pathways at high latitudes in the epipelagic layer, accounting for 0.50 in the Arctic Ocean and 0.40 in the Southern Ocean. This contribution decreases in temperate regions (0.30) and reaches its lowest values (0.10) in the oligotrophic subtropical gyres of the Pacific, Atlantic and Indian Oceans (**Fig. 3**). This gradient suggests a stronger ecological role for DNRA in colder, epipelagic, nutrient-rich environments, potentially reflecting enhanced nitrogen in these regions for energetic requirements.

### 3.3. Distribution of bioavailable nitrogen transformation pathways

We next examine the spatial distribution of nitrogen metabolic pathways involved in the transformation of bioavailable nitrogen (**Fig. 4**). We show that pathways driven by the metabolic requirement of assimilating nitrogen for cellular biosynthesis present distinct patterns from the ones using it as an electron acceptor for energy metabolism. ANRA, associated to cellular biosynthesis, exhibits its highest genomic potential in the subtropical gyres, with values exceeding 0.8 in the epipelagic layer (**Fig. 4a**, left map) and 0.5 in the mesopelagic layer (**Fig. 4a**, right map). Both latitudinal profiles show clear peaks around 30°N and 30°S, with additional seasonal variation at higher latitudes (∼60°N and 60°S) in the mesopelagic, where values range between 0.10 and 0.50 (**Fig. 4a**, right profile). In contrast, pathways related to nitrogen use for energy metabolism, such as nitrification and DNRA, show opposite spatial patterns to ANRA. Nitrification, associated with autotrophic processes in oxic environments, reaches its highest genomic potential at mid- to high latitudes (40–60°N and °S) in both depth layers (**Fig. 4b**). In the epipelagic Southern Ocean, however, projections are highly uncertain (NSD > 0.5; see **Fig. S3** for detailed uncertainty pattern). At high latitudes, nitrification potential in the epipelagic layer ranges from 0.15 to 0.30 between summer and winter (**Fig. 4b**, left panels), while in the mesopelagic it ranges from 0.40 to 0.80 between winter and summer (**Fig. 4b**, right panels). DNRA, typically associated with heterotrophic activity in suboxic to anoxic environments, shows peak genomic potential (∼0.5) in the high latitudes (∼60°N and °S) and, to a lesser extent, in equatorial upwelling regions (∼0.3), with relatively low seasonality in the epipelagic layer (**Fig. 4c**, left panels). In the mesopelagic layer, the genomic potential for DNRA is highest in equatorial upwellings and in the North Pacific (∼0.25; **Fig. 4c**, right panels). Additional hotspots are predicted in ODZ of the northern Indian Ocean and in the North Equatorial Countercurrent, with a high uncertainty however (NSD > 0.5; see **Fig. S3** for detailed uncertainty pattern). In conclusion, the spatial distributions of nitrogen transformation pathways associated to bioavailable forms of nitrogen reflect clear functional partitioning across ocean biomes and depth layers, shaped by two distinct metabolic requirements.

**Figure 4:**
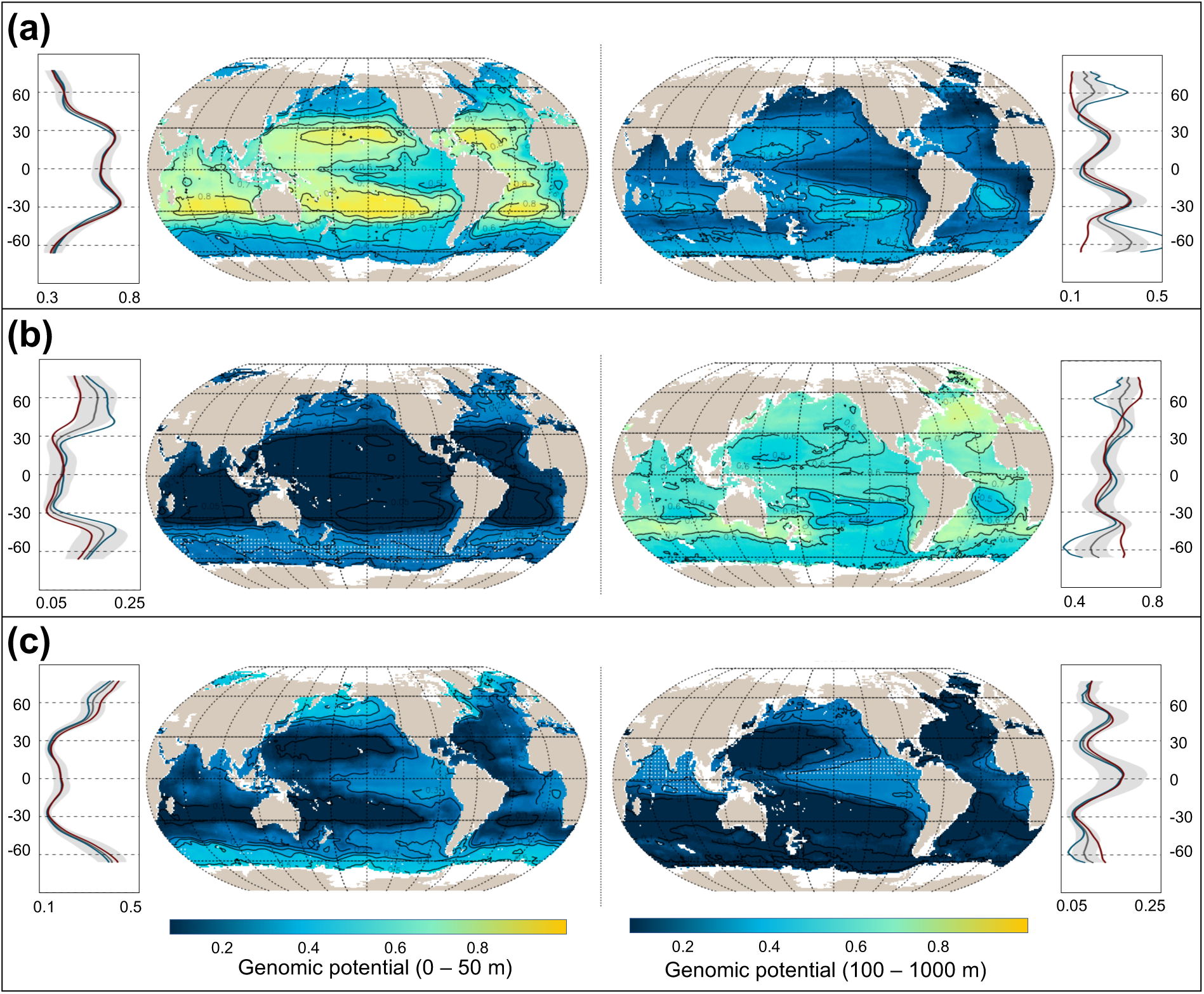
Genomic potential for bioavailable nitrogen transformation pathways represented by their relative abundance of metagenomic reads. Nitrogen transformation pathways are displayed in rows with: (a) Assimilatory Nitrate Reduction to Ammonium (ANRA), (b) Nitrification and (c) Dissimilatory Nitrate Reduction to Ammonium (DNRA). Left panels correspond to the epipelagic (0 – 50 m) projections and right panel correspond to the mesopelagic (100 – 1000 m) projections. Each panel represents the annual average spatial projection, with white stippling representing geographical areas associated with a high degree of uncertainty (i.e., standard deviation above 0.5 the global average). Latitudinal profiles are displayed in form of annual average (black), summer (red; northern hemisphere June to August) and winter (blue; northern hemisphere December to February). The projection uncertainty across bootstraps is presented as grey shading in the latitudinal profiles.

### 3.4. Distribution of bio-unavailable nitrogen transformation pathways

Nitrogen transformation pathways involving bio-unavailable forms (i.e., nitrogen fixation and denitrification) are less commonly encoded in microbial metagenomes than pathways associated with bioavailable nitrogen (**Fig. 3**). Here, we assess whether metabolic requirements structure their distribution similarly to pathways associated with bioavailable forms of nitrogen (**Fig. 5**). Nitrogen fixation, linked to the assimilation of bio-unavailable nitrogen for biosynthetic purposes, shows its highest genomic potential in the subtropical gyres, with values exceeding 0.7 in the epipelagic layer (**Fig. 5a**, left map) and 0.4 in the mesopelagic layer (**Fig. 5a**, right map). In the mesopelagic layer, however, projections are highly uncertain (NSD > 0.5; see **Fig. S3** for detailed uncertainty pattern). Latitudinal profiles show clear peaks around 30°N and 30°S, with moderate seasonality in the southern oligotrophic gyres of the epipelagic layer, where values range from 0.50 to 0.60 (**Fig. 5a**, right profile). In contrast, denitrification, associated with the transformation of ammonium to bio-unavailable nitrogen for energy metabolism, exhibits its highest genomic potential in high latitudes, tropical equatorial upwellings, and ODZ (**Fig. 5b**). In the epipelagic layer, values exceed 0.8 in high latitudes and 0.6 in equatorial upwellings. In the mesopelagic layer, characterized by lower dissolved oxygen, the genomic potential reaches values above 0.9 in high latitudes, tropical upwellings, and the Arabian Sea ODZ. In conclusion, metabolic requirements similarly shape the spatial distribution of nitrogen transformation pathways for both bio-unavailable and bioavailable forms.

**Figure 5:**
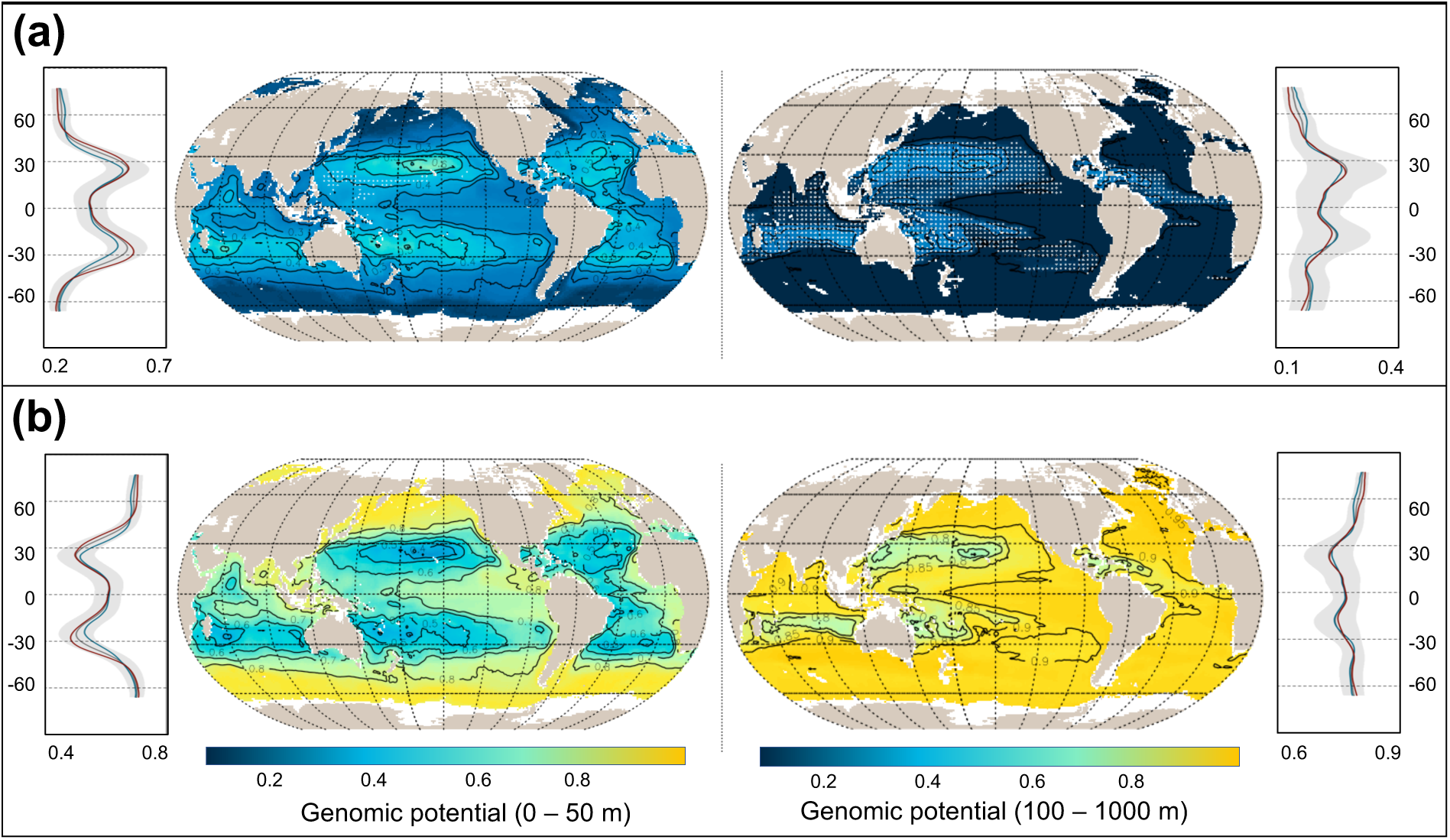
Genomic potential for bio-unavailable nitrogen transformation pathways represented by their relative abundance of metagenomic reads. Nitrogen transformation pathways are displayed in rows with: (a) Nitrogen fixation and (b) Denitrification. Left panels correspond to the epipelagic (0 – 50 m) projections and right panel correspond to the mesopelagic (100 – 1000 m) projections. Each panel represents the annual average spatial projection, with white stippling representing geographical areas associated with a high degree of uncertainty (i.e., standard deviation above 0.5 the global average). Latitudinal profiles are displayed in form of annual average (black), summer (red; northern hemisphere June to August) and winter (blue; northern hemisphere December to February). The projection uncertainty across bootstraps is presented as grey shading in the latitudinal profiles.

### 3.5. Environmental and taxonomic partitioning of nitrogen transformations

Nitrogen transformation pathways are tightly coupled to specific environmental and biogeochemical conditions, reflecting the distinct metabolic requirements of associated microbial communities, whether for cellular biosynthesis or energy metabolism (**Fig. 1**). To investigate the drivers of this partitioning, we identify the environmental features best explaining nitrogen transformation pathway composition, their overlap with environmental gradients, and the microbial communities supporting each pathway (**Text S1**, **Fig. 6**).

**Figure 6.**
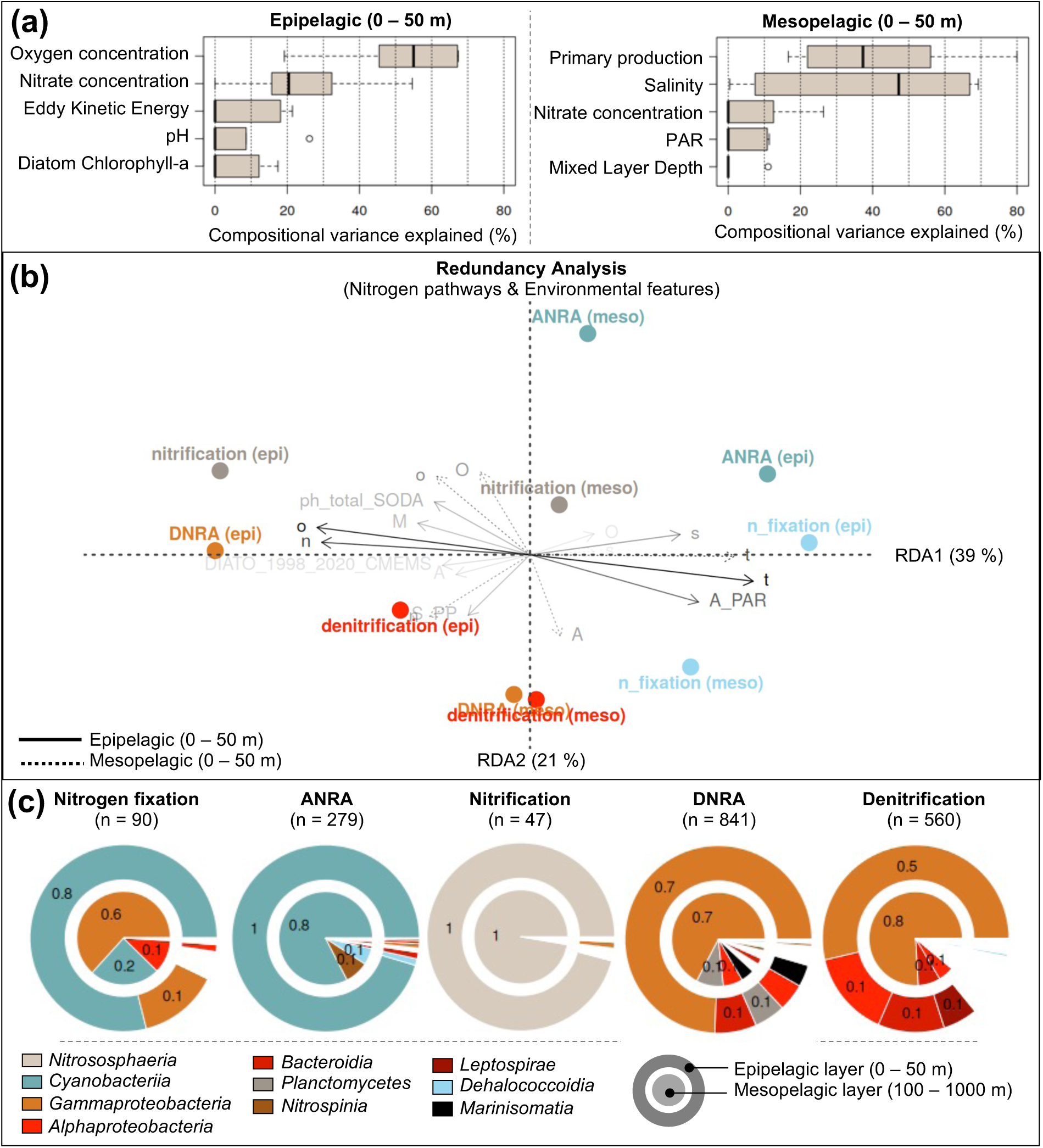
Environmental and taxonomic drivers of nitrogen transformation pathways. (a) Key environmental variables explaining pathway composition variance in the *in-situ* observations. (b) Redundancy Analysis (RDA) showing the relationship between nitrogen pathway partitioning and environmental composition; arrow transparency indicates the quality of representation on the first two RDA axes. (c) Key taxonomic classes supporting each nitrogen transformation pathway; n indicate the number of Metagenome-Assembled Genomes (MAGs) per pathway; epipelagic and mesopelagic layers are shown in the outer and inner pie charts, respectively.

In the epipelagic layer (**Fig. 6a**, left panel), pathway composition is primarily explained by oxygen (*median* 55% [*Q25–Q75*: 46–76%]) and nitrate concentrations (21% [16–32%]), with smaller contributions from eddy kinetic energy (0% [0–18%]), pH (0% [0–9%]), and diatom chlorophyll-*a* (0% [0–12%]). At the global scale, oxygen correlates with nitrate (ρ = 0.67) and acts as a proxy for temperature (–0.96), PAR (–0.63), and salinity (–0.51). In the mesopelagic model (**Fig. 6a**, right panel), surface primary production (37% [21–55%]) and salinity (47% [8–67%]) are the main features explaining the observed variance, with lower contributions from nitrate (0% [0–12%]) and PAR (0% [0–11%]). At the global scale, salinity correlates with temperature (0.65) and, to a lesser extent, nitrate (–0.43) and PAR (0.32). Therefore, oxygen and nitrate concentrations, temperature, and primary production, are the dominant environmental features structuring nitrogen transformation pathways across depth layers.

Distribution differences in nitrogen transformations related to biosynthetic versus energetic metabolic requirements is captured by the redundancy analysis (RDA), where the first (39%) and second (21%) axes represent the main gradients in the epipelagic and mesopelagic layers, respectively (**Fig. 6b**). In the epipelagic layer, RDA1 positively correlates with ANRA (r = 0.38), nitrogen fixation (0.45), temperature (0.91), and PAR (0.69), and negatively with nitrification (–0.50), DNRA (–0.51), denitrification (–0.21), nitrate (–0.85), and oxygen (–0.87). This axis links biosynthesis-related pathways to oligotrophic tropical environments and energy-related pathways to colder, nutrient-rich regions of the epipelagic ocean. The association between DNRA and oxic conditions may reflect a resolution artifact, missing micro-anoxic zones at high latitudes (**Fig. 6b**, and **4c**). In the mesopelagic layer, RDA2 correlates positively with ANRA (0.63) and oxygen saturation (0.82), and negatively with DNRA (–0.40), denitrification (–0.15), and oxygen utilization (–0.81), suggesting a partitioning between oxic and sub- or anoxic environments. Finally, mesopelagic nitrification shows weak correlations with both axes (0.04 and 0.14), consistent with its widespread dominance across the mesopelagic zone (**Fig. 6b** and **4b**).

Nitrogen transformation pathways are also supported by different communities between those associated to cellular biosynthesis, and energy requirements in oxic and sub- or anoxic environments (**Fig. 6c**). In the epipelagic layer, pathways associated with cellular biosynthesis and oxic conditions are primarily supported by autotrophic and aerobic bacteria such as *Cyanobacteria* (relative contribution between 0.8 and 1.0), with *Gammaproteobacteria* contributing to nitrogen fixation (0.1; **Fig. 6c**). Energy-related pathways under oxic conditions, such as nitrification, are predominantly associated with chemotroph archaea such as *Nitrososphaeria*. In contrast, pathways functioning under sub- or anoxic conditions are mainly supported by *Gammaproteobacteria* (0.5 to 0.7), the most genus-rich prokaryote class associated to obligatory or facultative anaerobes. We also observe lesser contributions from *Bacteroidia* (0.1), *Alphaproteobacteria* (0.1), *Leptospirae* (0.1 for denitrification), and *Planctomycetes* (0.1 for DNRA). In the mesopelagic layer, we show a similar taxonomic distribution as in the epipelagic layer for all nitrogen transformation pathways except for nitrogen fixation (**Fig. 6c**). In the later, non-cyanobacterial diazotrophs dominate the taxonomic composition, notably *Gammaproteobacteria* (0.6; **Fig. 6c**, left pie). We therefore highlight that metabolic requirement for nitrogen transformations partition, not only their distribution or environmental conditions they are associated with, but also the microbial communities supporting them.

## 4. Discussion

In this study, we examined whether metagenomics could capture the functional partitioning of nitrogen transformation pathways in the global ocean microbiome. We show that pathways linked to cellular biosynthesis and those tied to energy metabolism exhibit opposing spatial patterns, with biosynthetic pathways dominating in oligotrophic tropical gyres, while energy-related pathways peak in ODZs or high-latitude regions (**Fig. 4-5**).

### 4.1. Nitrogen transformation associated to cellular biosynthesis

Assimilatory processes such as ANRA and nitrogen fixation provide key functional advantages to autotrophic microbes in nutrient-depleted environments by supporting primary production under extreme oligotrophy (Herrero et al., 2019). These conditions prevail in subtropical gyres, where strong thermal stratification restricts vertical nutrient supply and maintains persistently low nitrate concentrations in surface waters (Dai et al., 2023). We showed elevated abundance of ANRA and nitrogen fixation encoding-genes in the epipelagic layer of tropical and subtropical regions, strongly associated with high temperature and low nitrate concentrations, consistent with known environmental controls on nitrate assimilation and nitrogen fixation (Luo et al., 2014; Zhang et al., 2020). The observed negative correlation with dissolved oxygen likely reflects its inverse relationship with temperature, as oxygen concentrations in surface waters remain far above thresholds limiting aerobic metabolism (>20 μmol kg⁻¹)(Dalsgaard et al., 2014; Lam and Kuypers, 2011). Metagenomics also reveals that both ANRA and nitrogen fixation are predominantly encoded by *Cyanobacteria*, including free-living phototrophs and symbiotic taxa capable of oxygenic photosynthesis and carbon fixation (Zehr, 2011). For example, the nitrogen-fixing cyanobacterium *UCYN-A* forms associations with photoautotrophic diatoms, relying on high PAR for energy (Farnelid et al., 2016; Zehr and Capone, 2020). Although we do not resolve species-level metagenomics, the association between nitrogen fixation encoding genes, *Cyanobacteria* abundance, and PAR in the epipelagic layer support this ecological link (**Fig.s 5a** and **6b** and **c**). These trends are further corroborated by the decline in genomic potential for both ANRA and nitrogen fixation with increasing latitude and depth, where light availability decreases (see environmental feature collection in Schickele et al., 2025). Finally, the dominance of ANRA over nitrogen fixation in terms of genomic potential (0.59 and 0.25 to 0.01 in the epi- and mesopelagic layers; **Fig. 4**) and number of supporting taxa (279 to 90; **Fig. 6c**) mirrors respective metabolic pathway rates (Gruber, 2008; Wang et al., 2019). Indeed, ANRA is widespread among autotrophic microbes, including diverse phytoplankton taxa (Gruber, 2008; Zehr and Kudela, 2011), whereas nitrogen fixation is restricted to a narrower and more specialized group (Messer et al., 2016; Turk-Kubo et al., 2023; Zehr, 2011).

### 4.2. Nitrogen transformation associated to energy requirements

Nitrogen transformations required for microbial energy metabolism also show strong consistency between metagenomic gene abundances and established biogeochemical knowledge (Battaglia and Joos, 2018; Lam and Kuypers, 2011; Zhang et al., 2023). Under oxic conditions, Nitrification encoding genes abundance, predominantly associated to *Nitrososphaeria* archaea (Ward, 2008), negatively correlates to PAR and positively correlates to nitrate concentration in the epi- and mesopelagic layers (Ward, 2008; Zhang et al., 2020). This matches the observed photoinhibition of ammonia oxidation in epipelagic waters (45–60 E m⁻² d⁻¹) versus its enhanced genomic potential in low-PAR high-latitudes and mesopelagic layers (2–25 E m⁻² d⁻¹)(see environmental feature collection in Schickele et al., 2025). Moreover, elevated nitrification gene abundance in high-nutrient, low-chlorophyll (HNLC) regions such as the North Pacific and Southern Ocean associates with ammonium accumulation driven by weak carbon export (Grundle et al., 2013; Sarmiento and Gruber, 2006). Therefore, the genomic potential of nitrification aligns with known nitrification distribution and rates (Pajares and Ramos, 2019).

The genomic potential for anaerobic nitrogen transformation pathways, such as denitrification and DNRA, positively correlates with low oxygen saturation and high apparent oxygen utilization (**Fig. 6b**). This is consistent with the dependence of *Gammaproteobacteria* on sub- or anoxic conditions to perform anaerobic respiration (Lam and Kuypers, 2011). The eastern Pacific upwelling, Arabian Sea, and Bay of Bengal ODZs show the highest genomic potential for DNRA and denitrification, reflecting known hotspots of anaerobic nitrogen transformation (Zhang et al., 2023). Interestingly, we detected a DNRA genomic potential mode at high latitudes in the epipelagic layer. These patterns are present in the *in-situ* observations (**Fig. 2b**) and associated by our model to large-scale oxygen climatologies (**Fig. 6b**). However, they may reflect the presence of micro-anoxic niches (e.g., within sea ice, particles, or sediments) or episodic sub- or anoxic conditions, such as freshwater influxes in the Arctic. Moreover, the genomic potential for DNRA in the epipelagic layer follows known distribution of Diatoms (**Fig. 6b**), that are known for their high degree of association with prokaryotes, and their use of DNRA for respiration under dark and anoxic conditions, such as high latitudes (Bellas et al., 2023; Kamp et al., 2011, 2015; Stief et al., 2022). In these cases, a genomic potential mode may not indicate active expression, but rather the retention of anaerobic potential in the genomes of facultative taxa. Indeed, many *Gammaproteobacteria* are facultative anaerobes, thus retain the genomic potential for anaerobic metabolism under oxic conditions (Pajares and Ramos, 2019). We also estimated a genomic potential for surface denitrification in the equatorial Pacific upwelling system, where gene expression (norB, nosZ) was already observed (i.e., although the exact mechanism is still unknown; Ganesh et al., 2015). However, this signal may also be amplified by vertical mixing, that episodically brings low-oxygen waters into the epipelagic zone. In more strongly stratified regions like the Arabian Sea and Bay of Bengal, where ODZs remain isolated from the epipelagic layer, we do not observe such a surface genomic potential, supporting the view that a fraction of these genes may be present in the surface microbiome but only actively expressed under sub- or anoxic conditions deeper in the water column (Dalsgaard et al., 2014; Zhang et al., 2023).

### 4.3. Metagenomic observations in biogeochemical modelling

We showed that the genomic potential for nitrogen transformation pathways reliably mirror associated biogeochemical patterns established by *in-situ* measurements and models (DeVries et al., 2012; e.g., Lam and Kuypers, 2011; Wang et al., 2019). Thus, metagenomics seems to be an observation-driven alternative to infer microbial nitrogen transformation patterns across diverse oceanic environments. While traditional biogeochemical models rely on bulk environmental proxies and generalized microbial groups, often limiting spatial and functional resolution (Bianchi et al., 2018, 2023; Deutsch et al., 2007), metagenomics directly captures the genetic potential for nitrogen transformations, providing mechanistic insights into microbial community functions (Louca et al., 2016). Our findings of nitrification, denitrification, and DNRA gene distributions correspond closely with known oxygen and light controls (Lam and Kuypers, 2011), reinforcing metagenomics as a valid proxy for microbial activity. This can be further developed towards transcriptomics and effective expression of those considered enzymes, or mechanistic approaches using the associations between genomic potential and environmental drivers (e.g., Salazar et al., 2019). Crucially, metagenomics reveals cryptic anaerobic niches and facultative metabolic capacities in ostensibly oxic zones and high latitudes. This micro-scale heterogeneity underscore metagenomics’ potential to capture episodic or fine-scale biogeochemical processes, although not completely understood yet. Moreover, integrating metagenomics with genome-scale metabolic models, such as those developed by Regimbeau et al. (2022) and Levine et al. (2025), offers a promising avenue to link gene content with microbial metabolic fluxes. These metabolic models leverage genomic data to predict microbial nutrient transformations under varying environmental conditions, thereby enhancing the predictive capacity of biogeochemical cycles (Levine et al., 2025; Régimbeau et al., 2022). However, they remain computationally demanding and there are very few existing metabolic models for marine organisms, in part because the organism must be easily cultured for biological validation (Levine et al., 2025). Together, these approaches enable the assimilation of cutting-edge *in-situ* observations into biogeochemical models, advancing our ability to predict microbial contributions to the ocean nitrogen cycle in a rapidly changing climate (Louca et al., 2016).

### 4.4. Caveats and limitations

Our approach is based on genomic potential, which captures the metabolic capabilities encoded in prokaryotic communities. However, it does not directly reflect metabolic rates or enzymatic activity, as it excludes gene expression and protein synthesis (Dalsgaard et al., 2014; Salazar et al., 2019). This limitation can be partially addressed through metatranscriptomic data, which provides insight into gene expression (Dalsgaard et al., 2014). However, metatranscriptomic exhibit a high spatial and temporal heterogeneity and are currently limited in sampling coverage (Paoli et al., 2022). Therefore, effort in both *in-situ* observations density and model resolution are needed to effectively capture the heterogeneity of gene expression in the global ocean. An additional limitation concerns the incomplete representation of certain nitrogen transformation pathways in existing gene databases. For instance, Anammox, an anaerobic process responsible for nitrogen removal in marine systems, could not be analysed here as only one specific gene was detected in our dataset (Paoli et al., 2022). Thus, the genomic potential for anammox could not be compared to other anaerobic processes like DNRA or denitrification (Eugster and Gruber, 2012; Lam and Kuypers, 2011; Yang et al., 2017). However, Anammox is estimated to contribute roughly one order of magnitude less to global nitrogen loss than DNRA or denitrification (Deng et al., 2024), suggesting that our analysis still captures the dominant nitrogen transformation pathways. Finally, two environmental variables known to influence nitrogen transformation pathways, iron, and ammonium concentration, were not included in our analysis. Global scale iron concentration remains difficult to represent at climatological scales, due to observational challenge, especially bioavailable iron, and model disagreements (Dai et al., 2023; Sohm et al., 2011). Compared to Nitrate, Ammonium concentration is known for its rapid turnover (0.05 against 370 years) and patchy distribution (Gruber, 2008), limiting its suitability for inclusion in climatological resolution models such as ours.

## 5. Conclusion

In this study, we show that the marine microbial metagenome captures distinct spatial patterns in nitrogen transformations associated with either cellular biosynthesis or energy metabolism. These patterns associate to distinct environmental drivers and microbial communities. Anaerobic and light-inhibited pathways, such as denitrification, nitrification, and DNRA, are enriched in eastern boundary upwelling systems, high-latitude regions, and deeper ocean layers. In contrast, genomic potential for ANRA and nitrogen fixation is concentrated in tropical oligotrophic gyres and surface ocean layers. This is consistent with known prokaryotic metabolic strategies and biogeochemical rates. Therefore, we suggest that metagenomics can complement or even substitute conventional biogeochemical measurements by offering spatially resolved, mechanistic insight into microbial nitrogen transformation pathways. This positions metagenomic data as a promising basis for new biogeochemical Essential Ocean Variables (EOVs; Muller-Karger et al., 2018). Beyond nitrogen, this framework may be extended to other elemental cycles, such as sulfur and methane metabolism (Salazar et al., 2019). Finally, to move from climatologies to near-real-time observations of microbial function, we emphasize the need for expanded *in-situ* metatranscriptomic sampling. The latter captures the temporal and spatial variability of gene expression, crucial to assess the response of microbial communities to short-term disturbances (e.g., marine heatwaves, oxygen minima) and long-term climate trends, ultimately advancing observation-based ocean biogeochemistry monitoring.

## Authors contribution

M.V., L.G., J-O.I. conceived and supervised the study and provided fundings. A.S. and H.S. processed the input data and performed the analysis. A.S., H.S. and N.G. wrote the first draft. A.S. conceived the modelling pipeline. All authors substantially contributed to the successive versions of this manuscript.

## Disclaimer

Copernicus Publications remains neutral with regard to jurisdictional claims made in the text, published maps, institutional affiliations, or any other geographical representation in this paper. While Copernicus Publications makes every effort to include appropriate place names, the final responsibility lies with the authors. Views expressed in the text are those of the authors and do not necessarily reflect the views of the publisher.

## Acknowledgements

This project has received funding from the European Union’s Horizon 2020 Research and Innovation Programme under grant agreement no. 862409 (Blue-Cloud), grant agreement no. 101094227 (Bluecloud2026) and no. 101059915 (BIOcean5D). This output reflects only the author’s view, and the European Union cannot be held responsible for any use that may be made of the information contained therein. The authors want to thank all the people involved in the Ocean Microbiomics database initiative and related scientific cruises for making their data publicly available. A.S. and H.S. would like to thank J. Härri, V. Sonnet, F. Benedetti, D. Eriksson, and C. Clerc for fruitful discussion on preliminary results and nitrogen transformation metabolic pathways.

## Code and data availability

All model outputs and R scripts used for data preparation, model analysis and figure generation are openly available at https://doi.org/10.5281/zenodo.18936759, under the Creative Commons Attribution 4.0 International license.

