## Supplementary for "Global pattern of nitrogen metabolism in marine prokaryotes"

### Text S1: Output pre-processing and statistical analysis

#### 1. Variance explained by environmental features

To identify key environmental features that explain an important fraction of the variance in the observations, we assessed the feature importance at the nitrogen transformation pathway level, which is the level at which we trained the model. We computed feature importance as the frequency with which each environmental feature was selected for tree splits, weighted by the corresponding reduction in model loss (Schickele et al., 2024). This measure highlights the environmental features that most effectively discriminate the composition of nitrogen transformation pathways and their relative abundance at the genome level, at each geographical location across the global ocean.

#### 2. Redundancy Analysis between Nitrogen pathways and environmental features

To examine how environmental conditions structure the distribution of nitrogen transformation pathways in the global ocean, we performed a Redundancy Analysis (RDA; Legendre & Legendre, 2012) including both the epi- and mesopelagic layers. Prior to analysis, pathway-level genomic potentials were transformed using a centered log-ratio (CLR; Aitchison, 1982) transformation for each depth layer, to account for the compositional nature of the data. We considered the five environmental features explaining most variance in either the epi- or mesopelagic layers. Moreover, we systematically considered sea water temperature, known as a major structuring parameter in the global ocean, even if not selected by CEPHALOPOD due to multicollinearity at the sampling locations (**Fig. S1**). All selected environmental features were integrated, when possible, over both the epi- and mesopelagic layers.

### 3. Microbial community associated to each pathway

To identify the microbial communities supporting each nitrogen transformation pathway, we extracted the class-level taxonomic composition associated with each selected enzymes and weighted them by their relative abundance. We chose the taxonomic class level as it corresponds to functionally coherent groups, relevant to biogeochemical processes (Louca et al., 2016) and offers cross-study comparability (Delmont et al., 2022; Paoli et al., 2022). We identified the top 10 taxa that most strongly support nitrogen transformations, based on their relative abundance across all considered pathways, for both epi- and mesopelagic layers.

### 4. Nitrogen transformation pathways composition by biomes

To compare the distribution and relative dominance of the genomic potential for nitrogen transformation pathways across the global ocean, we aggregated spatial projections at the biome level. We used the open-ocean biomes defined by Fay and McKinley (2014), which delineate large-scale ocean regions based on biogeochemical function. To ensure each biome contained *in situ* metagenomic observations and allowed for basin-level distinctions, we (i) merged the ‘ice biome’ and the ‘subtropical seasonally stratified biome’ into a single ‘Southern Ocean’ biome, and (ii) split the ‘subtropical permanently stratified biome’ by ocean basin into Pacific, Atlantic, and Indian sectors (see Figure 1 in Fay & McKinley, 2014). Within each biome and depth layer, we then calculated the average genomic potential from the spatial projections of the five nitrogen transformation pathways: nitrogen fixation, ANRA, nitrification, DNRA, and denitrification.

### 5. Isolating bio-unavailable nitrogen pathways

Given the compositional nature of the data, nitrogen transformation pathways like denitrification and nitrogen fixation can be challenging to interpret (e.g., Wang et al., 2019), especially when compared to DNRA, ANRA, and nitrification, which are an order of magnitude higher in relative abundance (see stacked barplots in **Fig. 2b**). To address this, we isolated the projections for nitrogen fixation and denitrification at both depth levels and recalculated their genomic potential independently of DNRA, ANRA, and nitrification. To do so, we divided the relative metagenomic reads of nitrogen fixation and denitrification each by the sum of the relative abundances of both. This approach alleviates the influence of more abundant processes when interpreting their respective spatial projection.

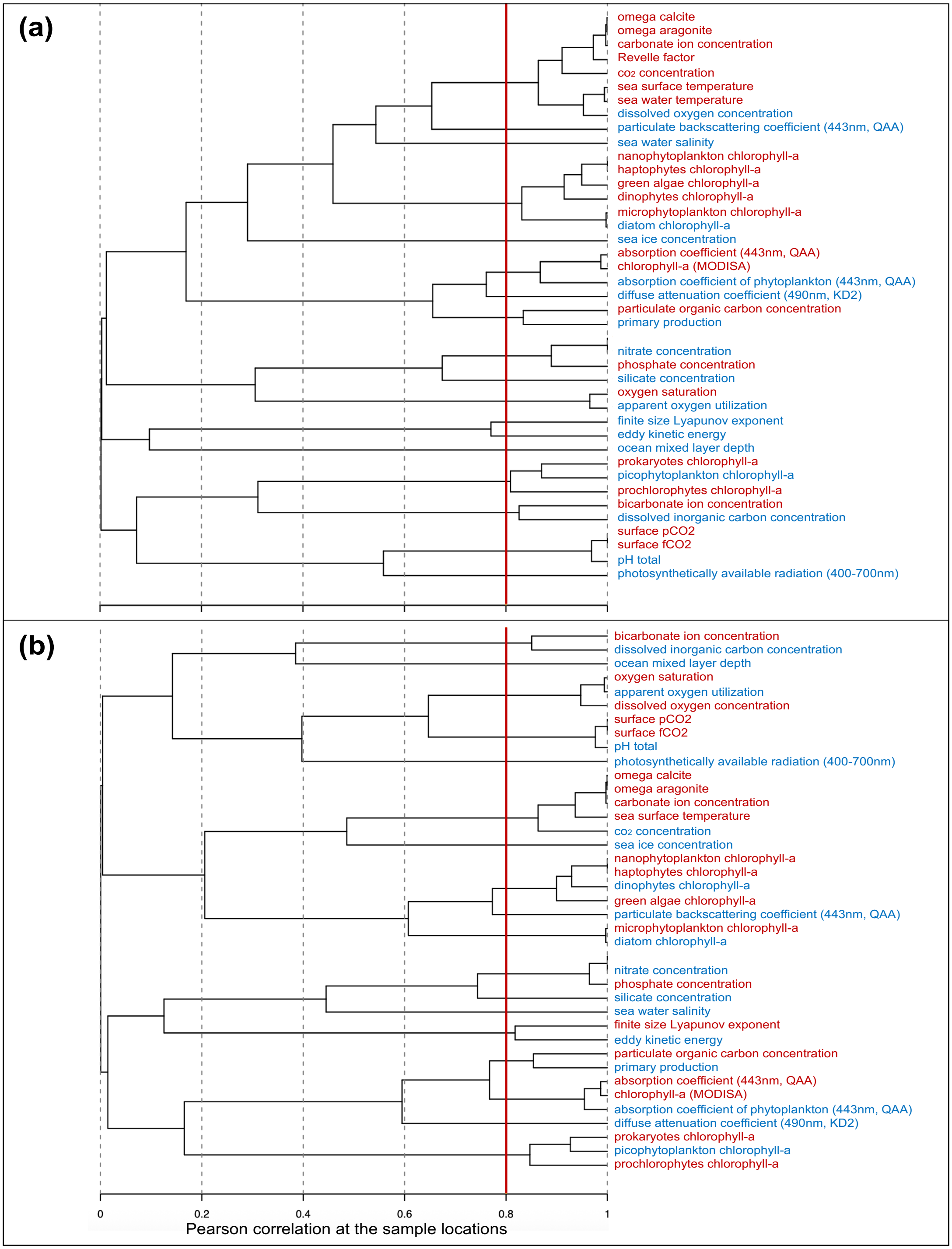


**Figure S1:** environmental feature correlation structure at the sample location for the (a) epi- and (b) mesopelagic layers. The considered environmental feature, within each cluster of intercorrelated features (Pearson > 0.8) are displayed in blue. Discarded features are displayed in red.

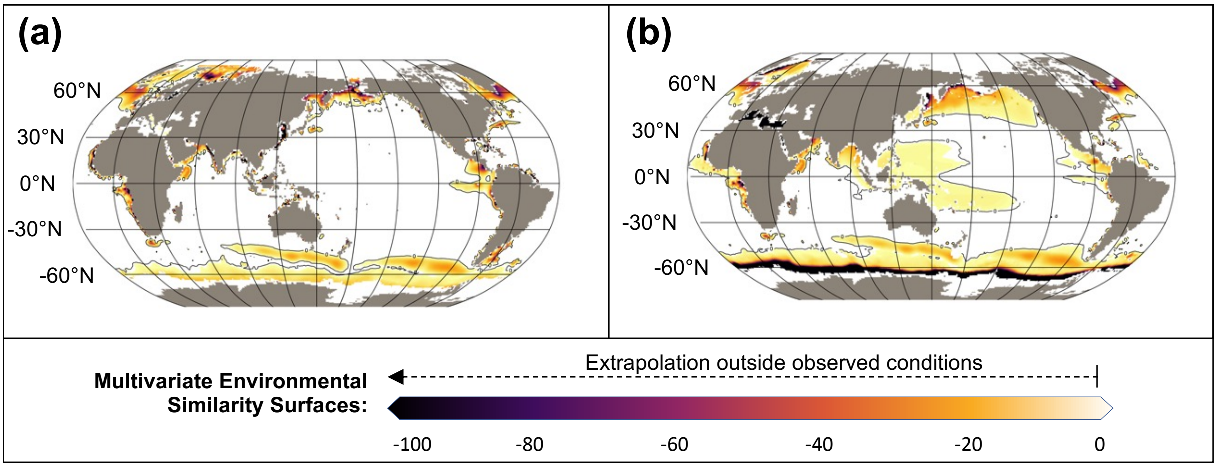


**Figure S2:** Multivariate Environmental Similarity Surface (MESS) analysis on the (a) epipelagic (0 – 50 m) and (b) mesopelagic (100 – 1000 m) observations. A negative value indicates extrapolation outside the environmental conditions characterizing the observations. The importance of the extrapolation is proportional to the MESS value.

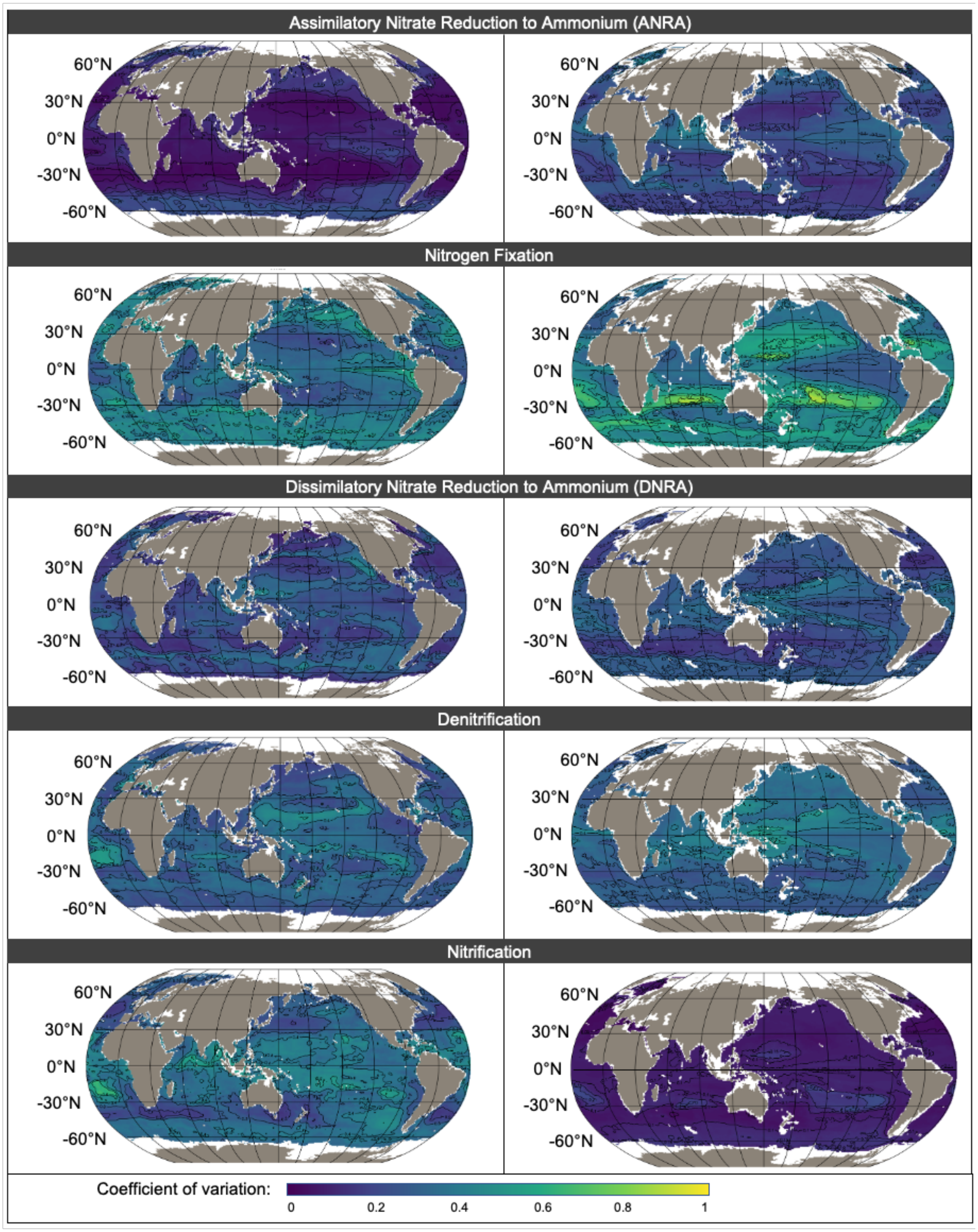


75 **Figure S3:** Coefficient of variation associated to the genomic potential for nitrogen transformation pathways represented by their relative abundance of metagenomic reads. Left panels correspond to the epipelagic (0 – 50 m) projections and right panel correspond to the mesopelagic (100 – 1000 m) projections. Nitrogen transformation pathways are displayed in rows with: Assimilatory Nitrate Reduction to Ammonium (ANRA), Nitrogen fixation, Dissimilatory Nitrate Reduction to Ammonium (DNRA), Denitrification, and Nitrification.

**Table S1**: Collection of observation-based, monthly-resolved environmental features at the global scale derived from WOA18 (Garcia et al. 2018), Ocean Soda (Gregor and Gruber, 2020), GMIS (NASA, 2024), CMEMS (Xi et al. 2021) and AVISO (D'Ovidio, 2022).

| Name | Unit | Source | Depth resolution | Time range |
|---|---|---|---|---|
| ocean mixed layer thickness | m | WOA18 | NA | 1981-2010 |
| apparent oxygen utilization | $\mu mol.kg^{-1}$ | WOA18 | 0 – 50 m and 100 – 1000 m | 1900-2017 |
| silicate concentration | $\mu mol.kg^{-1}$ | WOA18 | | 1900-2017 |
| nitrate concentration | $\mu mol.kg^{-1}$ | WOA18 | | 1900-2017 |
| dissolved oxygen concentration | $\mu mol.kg^{-1}$ | WOA18 | | 1900-2017 |
| oxygen saturation | % | WOA18 | | 1900-2017 |
| phosphate concentration | $\mu mol.kg^{-1}$ | WOA18 | | 1900-2017 |
| sea water salinity | - | WOA18 | | 1955-2017 |
| sea water temperature | °C | WOA18 | | 1955-2017 |
| co2 concentration | $\mu mol.kg^{-1}$ | OceanSoda | Surface only | 1982-2022 |
| carbonate ion concentration | $\mu mol.kg^{-1}$ | OceanSoda | | 1982-2022 |
| dissolved inorganic carbon concentration | $\mu mol.kg^{-1}$ | OceanSoda | | 1982-2022 |
| bicarbonate ion concentration | $\mu mol.kg^{-1}$ | OceanSoda | | 1982-2022 |
| sea ice concentration | - | OceanSoda | | 1982-2022 |
| omega aragonite | - | OceanSoda | | 1982-2022 |
| omega calcite | - | OceanSoda | | 1982-2022 |
| pH total | -log([H+]) | OceanSoda | | 1982-2022 |
| revelle factor | - | OceanSoda | | 1982-2022 |
| surface fCO2 | - | OceanSoda | | 1982-2022 |
| surface pCO2 | μatm | OceanSoda | | 1982-2022 |
| total alkalinity | $\mu mol.kg^{-1}$ | OceanSoda | | 1982-2022 |
| absorption coefficient (443nm, QAA) | $m^{-1}$ | GMIS | | 2002-2017 |
| absorption coefficient of phytoplankton (443nm, QAA) | $m^{-1}$ | GMIS | | 2002-2017 |
| total alkalinity (SVR ensemble) | $\mu mol.kg^{-1}$ | GMIS | | 2002-2017 |
| particulate backscattering coefficient (443nm, QAA) | $m^{-1}$ | GMIS | | 2002-2017 |
| chlorophyll-a (MODISA) | $mg\ Chl.m^{-3}$ | GMIS | | 2002-2017 |
| diffuse attenuation coefficient (490nm, KD2) | $m^{-1}$ | GMIS | | 2002-2017 |
| photosynthetically available radiation (400-700nm) | $Ein\ m^{-2}.d^{-1}$ | GMIS | | 2002-2017 |
| primary production | $mgC\ m^{-2}.d^{-1}$ | GMIS | | 1997-2010 |
| particulate organic carbon concentration | $mgC.m^{-3}$ | CMEMS | | 1998-2010 |
| diatom chlorophyll-a | $mgChl.m^{-3}$ | CMEMS | | 1998-2020 |
| dinophytes chlorophyll-a | $mgChl.m^{-3}$ | CMEMS | | 1998-2020 |
| picophytoplankton chlorophyll-a | $mgChl.m^{-3}$ | CMEMS | | 1998-2020 |
| prochlorophytes chlorophyll-a | $mgChl.m^{-3}$ | CMEMS | | 1998-2020 |
| prokaryotes chlorophyll-a | $mgChl.m^{-3}$ | CMEMS | | 1998-2020 |
| green algae chlorophyll-a | $mgChl.m^{-3}$ | CMEMS | | 1998-2020 |
| microphytoplankton chlorophyll-a | $mgChl.m^{-3}$ | CMEMS | | 1998-2020 |
| nanophytoplankton chlorophyll-a | $mgChl.m^{-3}$ | CMEMS | | 1998-2020 |
| haptophytes chlorophyll-a | $mgChl.m^{-3}$ | CMEMS | | 1998-2020 |
| eddy kinetic energy | $cm^2.s^{-2}$ | AVISO | | 1993-2021 |
| finite size Lyapunov exponent | $d^{-1}$ | AVISO | | 2000-2021 |
